# A simple island biodiversity model is robust to trait dependence in diversification and colonization rates

**DOI:** 10.1101/2022.01.01.474685

**Authors:** Shu Xie, Luis Valente, Rampal S. Etienne

## Abstract

The application of state-dependent speciation and extinction (SSE) models to phylogenetic trees has revealed an important role for traits in diversification. However, this role remains comparatively unexplored on islands, which can include multiple independent clades resulting from different colonization events. Here, we perform a robustness study to identify how trait-dependence in rates of island colonization, extinction and speciation (CES rates) affects the estimation accuracy of a phylogenetic model that assumes no rate variation between trait states. We extend the DAISIE (Dynamic Assembly of Islands through Speciation, Immigration and Extinction) simulation model to include state-dependent rates, and evaluate the robustness of the DAISIE inference model using simulated data. Our results show that when the CES rate differences between trait states are moderate, DAISIE shows negligible error for a variety of island diversity metrics. However, for large differences in speciation rates, we find large errors when reconstructing clade size variation and non-endemic species diversity through time. We conclude that for many biologically realistic scenarios with trait-dependent speciation and colonization, island diversity dynamics can be accurately estimated without the need to explicitly model trait dynamics. Nonetheless, our new simulation model may provide a useful tool for studying patterns of trait variation.

## Introduction

Evolutionary biologists have long studied how functional traits affect macroevolution (Stanely, 1975; Cracraft, 1985; Jablonski, 2008; Rabosky and McCune, 2010; Rabosky and McCune, 2010; Simpson, 2013; Chevin, 2016). A suite of empirical analyses have demonstrated that a diverse set of traits, such as body size (Mattila and Bokma, 2008; Lee et al., 2008), migratory behavior (Rolland et al. 2014), sexual conflict (Arnqvist et al., 2000), and diet (Farrell, 1998; Burin et al., 2016), have significant effects on diversification rates.

Revealing the role of traits in diversification requires statistical and modeling approaches. When a certain trait state occurs more frequently in a speciose clade than in a species-poor clade, we may be tempted to conclude that this state promotes speciation or reduces extinction. However, such a pattern may be due to the evolutionary conservation of the trait (Rabosky and Goldberg 2015). An excess of species with a particular state may also be due to asymmetrical transitions between states (Goldberg and Igić, 2012; Burin et al., 2016). The increasing availability of molecular phylogenies has stimulated the development of statistical methods to detect how traits are associated with diversification (Mitter et al., 1988). The earliest methods including sister clade comparisons could only address the variation in net diversification rates (speciation minus extinction), but could not distinguish the effect of a trait on speciation and extinction separately (Barraclough, 1998; Farrell, 1998; Heilbuth, 2000; Maddison et al., 2007). The binary-state-dependent speciation and extinction model (BiSSE), a likelihood-based framework, resolved these shortcomings (Maddison et al. 2007a). BiSSE inspired a large number of state-dependent diversification models, which are known as the SSE (state-dependent speciation and extinction) model family. These models extend BiSSE in various ways (e.g. considering more than two trait states, continuous rather than discrete states, spatial location as a trait) to enable state-dependent analyses to infer state-dependent diversification under a variety of scenarios or to model more complex phenotypic traits (Fitzjohn et al., 2009; Fitzjohn, 2010; Goldberg et al., 2011; Goldberg and Igić, 2012). Furthermore, models incorporating hidden traits were developed to overcome high type I errors with BiSSE (Beaulieu and O’Meara, 2016; Herrera-Alsina et al., 2019).

One geographical setting where traits have long been proposed to influence diversity are islands (Parent et al., 2008; Cowie and Holland, 2008). Oceanic islands are home to some of the most extraordinary radiations, such as Darwin’s finches (Losos and Ricklefs, 2009) and a key question is whether certain traits or trait states have played a role in the presence or absence of rapid radiations. While insular radiations are thought to be mostly driven by increased ecological opportunity on islands, it has long been hypothesized that some traits may trigger, facilitate or hinder diversification in an insular setting (García-Verdugo et al., 2014; Zhu et al., 2020). Some characteristics of species, such as seed size in plants and flight ability in birds, evidently affect the chances of species colonizing an island or an isolated habitat (Onstein et al., 2017). Furthermore, after successful colonization of an island, changes in morphological characteristics occur, facilitated by ecological release (Losos 1997; Millien, 2006). These character changes have been shown to affect *in situ* diversification rates of species, which are important to address evolutionary assembly on islands (Aleixandre et al., 2013; Burns, 2016; Biddick et al., 2019).

Despite the fundamental role of islands in the conceptual development of trait diversification theory, investigations of trait diversification dynamics on islands in a phylogenetic context are rare compared to large continental radiations. One reason for this is that insular communities are typically less diverse than continents, and thus their phylogenies are comparatively small and information-poor; therefore they are not amenable to fitting SSE models, which generally require relatively large phylogenetic trees (Davis et al., 2013). Thus, most insular radiations cannot be studied using classic SSE approaches. Alternatively, one can study multiple phylogenetic trees simultaneously. While SSE models were not designed for this purpose, it is relatively straightforward to extend them to apply to multiple trees. However, insular communities assemble via colonization and potentially subsequent diversification, and thus focusing only on trees of clades that have radiated (and not on colonization times or singleton lineages that have not diversified) ignores an important part of the processes that form insular communities.

Current inference methods for studying island community assembly and diversification ignore rate variation between species with different trait states. For example, the dynamic stochastic island biogeography model DAISIE (Dynamic Assembly of Islands through Speciation, Immigration and Extinction) (Valente et al., 2015, 2018, 2020) allows estimation of colonization and diversification rates from the colonization and branching times of insular communities, which can be obtained from the collection of multiple phylogenetic trees resulting from several colonizations of an island (e.g. all mammals on an island). However, the DAISIE framework is currently silent on the effects of traits of insular species, which may lead to erroneous estimates of parameters (colonization, speciation and extinction) if species traits help shape the phylogenetic trees on islands. Currently, there is no likelihood-based inference model focusing on phylogenetic data from islands that incorporates both trait dynamics and their effects on diversification rates (an island version of an SSE model, or an SSE version of DAISIE) because deriving the likelihood for a model with both diversity-dependent and trait-dependent diversification is not trivial.

In the absence of a method to estimate trait state-dependent colonization, extinction and speciation rates (CES rates) from island communities, can we still obtain meaningful results regarding island diversification using current island biogeography models? Specifically, although DAISIE does not include trait dynamics, can it nevertheless accurately reconstruct island diversity dynamics through time? Every model is a simplification of reality, and leaves out details that are relatively irrelevant (Friedman 1953). In this spirit we ask: are trait dynamics irrelevant for understanding diversification dynamics? Under what conditions of trait dynamics can trait-less DAISIE still be used to make accurate predictions of island diversity, distribution of island clade sizes, and diversity changes through time? In this paper we present a pipeline to answer these questions. Specifically, we assess i) whether DAISIE is able to accurately reconstruct island species assemblage in the presence of asymmetric rates between binary trait states, ii) how the performance of the model is influenced by unequal rates of transition between trait states, and iii) which CES rate has the largest effect on the robustness of DAISIE. If we find that in the presence of trait dependence DAISIE can accurately infer island diversity dynamics through time even without explicitly modelling such trait dependence, this will suggest that simple models do a good job of explaining island diversity and that reliable analyses can be performed even without trait data of island species (which are often absent, incomplete, or difficult to obtain), i.e. the model is robust to trait dynamics. If, instead, we find that under certain conditions the existence of trait-dependence substantially alters the predictions of DAISIE, this will suggest that traits cannot be ignored in these cases, and that a new estimation approach is needed.

## Methods

### State-dependent and state-independent simulation models

The DAISIE inference framework uses maximum likelihood to estimate colonization and diversification rates of insular biota from phylogenetic information. The core version of DAISIE assumes that all island species share the same CES rates, and the model is essentially neutral at the species-level (Valente et al. 2015). However, dynamics of state-dependent diversification and colonization within island clades are not modelled.

Here, we introduce a trait state-dependent island biodiversity simulation model, an extension of the DAISIE simulation model combining it with features of the BiSSE model (Fig. 1). To distinguish the two simulation models, the new simulation model is termed *state-dependent simulation* (SDS) model, and the original trait-less DAISIE simulation model is termed *state-independent simulation* (SIS) model. Likewise, we will call the standard DAISIE inference model the *state-independent inference* (SII) model. A state-dependent inference (SDI) model does not yet exist. In the SDS model, the rates of all evolutionary processes are state-dependent. For simplicity, we consider a binary trait with two states, 1 and 2. Species in the same state have the same CES rates, while species with different states may differ in one or more rates. Mainland species can be regarded as forming two assemblages according to their trait states. Immigration of species in each assemblage to the island is determined by the number of mainland species in each state (*M*_1_ and *M*_2_) and their colonization rates (*γ_1_* and *γ_2_*). Once an immigrant species (which inherits the trait state from its mainland ancestor) successfully colonizes the island, it can undergo population divergence from the mainland population (via anagenesis *λ_1^a^_* and *λ_2^a^_*,), *in situ* speciation (via cladogenesis *λ_1^c^_* and *λ_2^c^_*) or extinction (*μ_1_* and *μ_2_*). Island species can shift between trait states at a certain rate of transition from state 1 to state 2 (*q_12_*) or from state 2 to state 1 (*q_21_*). The transition rates can be equal or different. In speciation via anagenesis or cladogenesis, daughter lineages are assumed to inherit the trait state of their parent species, which means no state shifts occur during speciation (FitzJohn, 2010). Furthermore, transitions are regarded as intraspecific changes, which occur instantaneously along lineages, thus assuming that the period of time in which two states coexist in a polymorphic species is negligible. Transition and speciation events are not allowed to occur simultaneously in this model (Maddison et al., 2007). The equations for calculating the rates are given in the Supplementary Methods.

**Fig 1:**
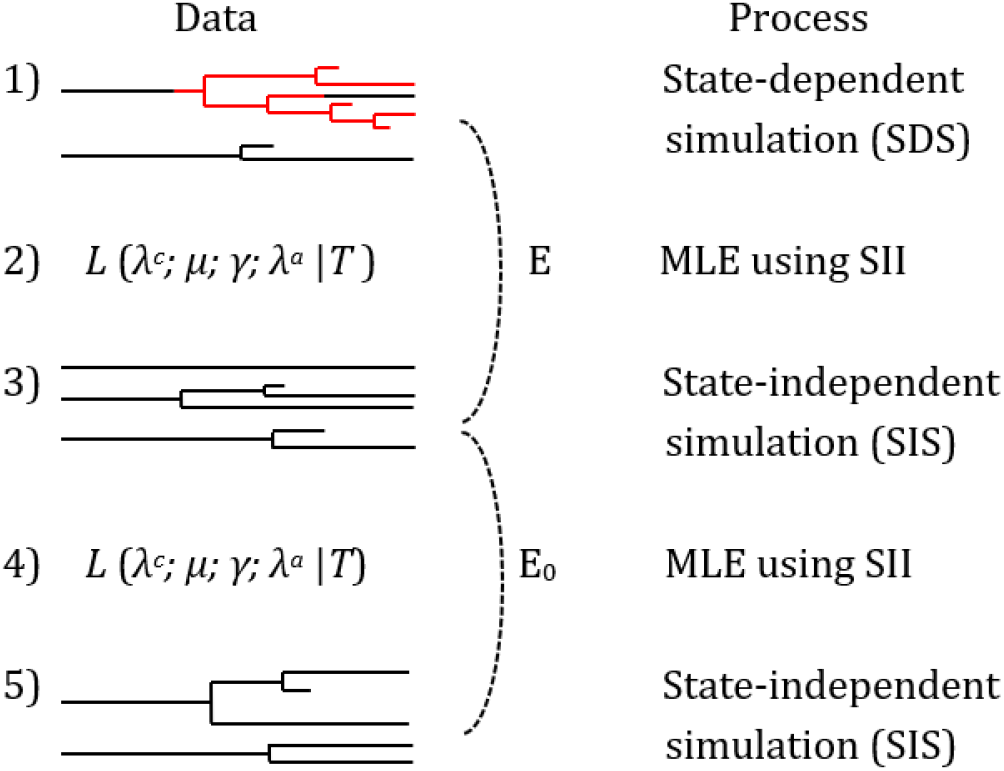
Schematic representation of the robustness pipeline. (1) Simulate phylogenetic data with the SDS model. The binary states are represented by two different colors (red and black). (2) Use the data obtained from step 1 to estimate parameters with the SII model. (3) Simulate data using the SIS model with parameters estimated in step 2. (4) Use the SII model again to estimate parameters. (5) Simulate data using the SIS model with the estimated parameters from step 4.

In both SIS and SDS simulations, we considered diversity-dependent (DD) and diversity-independent (DI) models. In DI models, all the rates are diversity-independent. In DD models, colonization and cladogenetic speciation rates are diversity-dependent, while the other rates are diversity-independent. We implemented a clade-specific DD model, assuming that diversity-dependence only operates between species in the same clade, which descend from the same mainland ancestor. We also assumed, for the sake of simplicity, that diversity-dependence is not state-dependent, i.e. the diversity-dependent term is the same regardless of the trait state of the species undergoing speciation or colonization. It is possible to implement the combination of diversity-dependence and trait-dependence in our simulation model; however, we do not have strong evidence to indicate it is necessary. We assumed that differences in resource utilization due to phylogenetic distance in species belonging to different clades are sufficient to prevent competition between clades. We implemented the SDS model in the R package DAISIE (Etienne et al., 2021).

The SDS results record the evolutionary history of island species including their colonization and branching times, as well as the richness dynamics of each trait state. We assumed that the inference accuracy of the DAISIE model will be poorer for larger inequality between the numbers of species in each of the two states. To test this assumption, we used the tip ratio *r* (Davis et al,. 2013), which we denoted by *r*, as the number of species in the species-rich state divided by the number of species in the species-poor state:

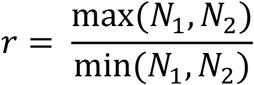

We used seven island diversity metrics to evaluate the simulated phylogenies from the SDS and the SIS models. Four metrics were used to measure diversity at the end of the simulation: total number of species (*N*_Spec_); number of lineages present on the island (*N*_Col_) (resulting from independent colonization events), standard deviation of clade size (σ_CS_) and standard deviation of colonization time among clades (σ_CT_). The other three metrics measured richness changes through time: total species richness through time (SRTT); endemic species richness through time (ESRTT); non-endemic species richness through time (NESRTT).

### Simulation scenarios

The mainland pool of 1000 species is assumed to be evenly distributed with 500 species in each state. There is no loss of generality depending on the mainland pool because it is the difference in total colonization rate (the product of mainland pool size and per capita colonization rate) that matters (Valente et al. 2015). In addition, we set a limit of 20 (*K’* = 20) species for each clade for the diversity-dependent model. To measure the effect of transition rates independent of CES rates, we used a symmetric scenario as a control, where all the CES rates are symmetric between binary states (Table 1A). We chose two values for each CES rate, a low one and a high one, in such a way that the total number of species remains between realistic values of 50 to 150. We set four types of transitions between binary states (Table 1B). For high and low transition rates we used 0.2 and 0.02, respectively. The symmetric scenario consists of 128 combinations of CES rates, transition rates and diversity dependence (Table 1A).

**Table 1.** Parameter space for the state-dependent simulations. **A.** The symmetric (control) scenario with all the CES rates symmetric between binary states, consisting of each combination of the parameters. Numbers separated by comma are the low and high values, respectively. **B.** Four transition types with different combinations of *q*_12_ and *q*_21_. **C.** The 13 scenarios of asymmetric CES rates (*λ^c^, λ^a^, γ, μ*). The control scenario contains the parameter sets where all the CES rates are symmetric (RRD = 0). For each asymmetric scenario, only one of the four rates is considered to be asymmetric (RRD ≠ 0). The degree of asymmetry is represented by the relative rate differential (RRD). Larger RRD value indicates larger rate difference between states.

| **A.** | **Parameters** | **Parameter values** |
| --- | --- | --- |
|  | Time | 5 |
|  | M1 | 500 |
|  | M2 | 500 |
| | $\lambda^c_1 = \lambda^c_2$ | 0.2, 0.4 |
| | $\mu_1 = \mu_2$ | 0.1, 0.2 |
| | $\gamma_1 = \gamma_2$ | 0.008, 0.012 |
| | $\lambda^a_1 = \lambda^a_2$ | 0.2, 0.4 |
| | $q_{12}$ | 0.02, 0.2 |
| | $q_{21}$ | 0.02, 0.2 |
| | $K'$ | 20, ∞ |

| **B.** | **Transition types** | **$q_{12}$** | **$q_{21}$** |
| --- | --- | --- | --- |
| | low $q_{12}$ low $q_{21}$ | 0.02 | 0.02 |
| | high $q_{12}$ low $q_{21}$ | 0.2 | 0.02 |
| | low $q_{12}$ high $q_{21}$ | 0.02 | 0.2 |
| | high $q_{12}$ high $q_{21}$ | 0.2 | 0.2 |

| **C.** | **Symmetry of rates** | **Scenario** | **RRD of cladogenesis ($\lambda^c$)** | **RRD of extinction ($\mu$)** | **RRD of colonization ($\gamma$)** | **RRD of anagenesis ($\lambda^a$)** |
| --- | --- | --- | --- | --- | --- | --- |
|  | Symmetry (control) | 1 | 0 | 0 | 0 | 0 |
|  | Asymmetry in cladogenesis | 2 | 0.5 | 0 | 0 | 0 |
|  |  | 3 | 1 | 0 | 0 | 0 |
|  |  | 4 | 1.5 | 0 | 0 | 0 |
|  | Asymmetry in extinction | 5 | 0 | 0.5 | 0 | 0 |
|  |  | 6 | 0 | 1 | 0 | 0 |
|  |  | 7 | 0 | 1.5 | 0 | 0 |
|  | Asymmetry in colonization | 8 | 0 | 0 | 0.5 | 0 |
|  |  | 9 | 0 | 0 | 1 | 0 |
|  |  | 10 | 0 | 0 | 1.5 | 0 |
|  | Asymmetry in anagenesis | 11 | 0 | 0 | 0 | 0.5 |
|  |  | 12 | 0 | 0 | 0 | 1 |
|  |  | 13 | 0 | 0 | 0 | 1.5 |

To investigate the effect of trait-dependence, we ran the SDS simulation under a series of scenarios with varying degrees of asymmetry in CES rates. In these scenarios, the mean values of CES rates between binary states were kept the same as the symmetric scenario, as well as two gradients of mean rates for each parameter (low and high, Table 1A). For the analyses with asymmetry in rates, only one CES rate was set to be asymmetric per scenario, and all the others were kept symmetric (Table 1C). We defined the relative rate differential (RRD) as the difference in rates for the two trait states by their mean rate value, to measure the asymmetry level between states.

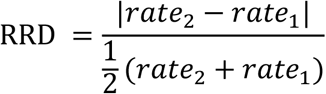

RRD is 0 when the rates are symmetric for the two states, and larger RRD means a larger rate difference between states. For each CES rate, we ran analyses with three different levels of RRD: 0.5, 1, 1.5 (Table 1C). Therefore, in total we ran 13 scenarios: 1 control plus 3 asymmetric scenarios for each of the 4 parameters (rates of cladogenesis, extinction, colonization and anagenesis) (Table 1C). We did not consider combinations of asymmetric rates as the number of parameter combinations would have been computationally prohibitive. We set CES rate values of state 2 to always be higher or equal to the values of state 1 for all the asymmetric cases to avoid redundancy. In asymmetric scenarios, transition rates were chosen in the same way as for the symmetric scenario, with four transition types (Table 1B).

### Robustness analysis

We aimed to test whether ignoring trait dynamics in inference affects the ability of the SII model to reconstruct diversity dynamics on islands. We used a robustness computational pipeline (Fig. 1) to measure the error of the SII model when real insular diversity dynamics involve trait dependence in rates (SDS), adapting the approach of Neves et al. (2021). Parameters were estimated on either SDS or SIS phylogenies using the SII in the pipeline. First, we simulated 1000 replicates (‘islands’) for each parameter set (Table 1), under the SDS model. We converted the SDS results to the SIS output format (i.e. stripped away of trait-related information), and estimated CES rates and carrying capacity with the SII model. We then used the estimated parameters to simulate with the SIS model, and again estimated the parameters for these simulations with the SII model and used the obtained parameters to simulate a second set of SIS results. The difference between the inferences on the two SIS results gives the baseline error (*E*_0_) in inference that occurs even if the inference model is identical to the simulation model (SII = SIS). The difference between the inferences on the SDS results and the first SIS results gives the error *E* that occurs when the simulation and inference model differ (SII ≠ SDS) (Fig. 1).

We calculated the errors *E* and *E_0_* of seven metrics for each replicate, which resulted in two error distributions for each parameter set. *E* and *E_0_* for *N*_Spec_, *N_Col_*, σ_CS_ and σ_CT_ were calculated as the absolute difference between simulations, while for the three diversity-through-time metrics (ΔSRTT, ΔESRTT, ΔNESRTT), errors were calculated using the ΔnLTT (normalized lineage-through-time) statistic (Janzen et al., 2015), by integrating the absolute distance between two diversity-through-time curves (Fig. S1). ΔnLTT is equal to zero only when the two simulated nLTT curves are identical. To compare the distributions of the two errors, *E* and *E*_0_, we used a metric, *ED_95_* (Neves et al., 2021). ED_95_ is the percentage of the distribution of *E* that exceeds the 95% percentile of the distribution of *E*_0_. Higher ED_95_ values indicate larger differences between *E* and *E*_0_. The ED_95_ of all the seven island diversity metrics were calculated for each parameter set. We ran a total of 1664 parameter sets (128 for each of the 13 scenarios), with 1000 replicates for each set. The calculation of metrics and error analysis were implemented in the R package DAISIErobustness (Neves et al. 2021).

## Results

We find that DAISIE is quite robust to trait-dynamics. The inference errors are negligible for all metrics except ΔNESRTT (non-endemic richness through time) and σ_CS_ (clade size standard deviation), and the latter are only affected when cladogenetic speciation rates are asymmetric. In addition, diversity-dependence and state-dependent transition rates have negligible effect on the inference errors except under asymmetric cladogenesis. Surprisingly, the inference errors are not related to the tip ratio, i.e. the ratio of the maximum and minimum diversities of each trait state, but they are positively correlated with the variation in clade sizes. We now present these results in more detail.

### State-dependent CES rates and the inference errors

In general, using the SII model to estimate parameters and subsequently the SIS model to simulate with the obtained parameters causes minimal error in reconstructing the SDS diversity patterns for most of the parameter sets (Fig. 2). Most metrics measuring the diversity at present (*N*_Spec_, *N*_Col_ and σ_CT_) show minimal difference between *E* and *E_0_* for all scenarios (Fig. 2). However, σ_CS_ shows the largest error difference among all the metrics, and is positively correlated to the RRD of the cladogenesis rate. Among the three ΔnLTT statistical metrics, only the error in ΔNESRTT is large when there is high asymmetry in cladogenesis rate (RRD = 1.5) (Fig. 2).

**Fig 2.**
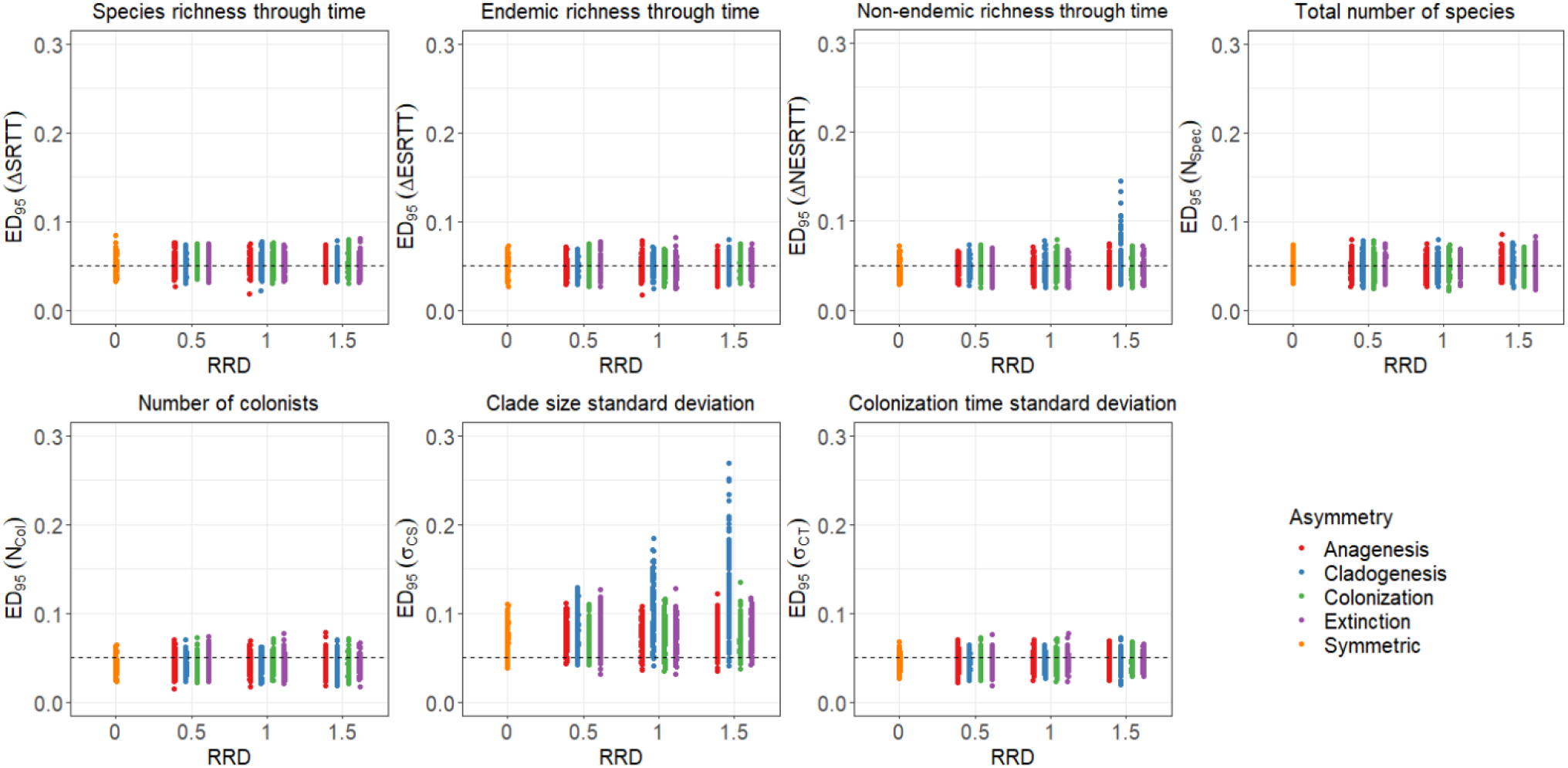
Distribution of *ED_95_* for seven island diversity metrics. Each point corresponds to a parameter set. The orange points correspond to the parameter sets in the control scenario with all CES rates symmetric, and the other colors correspond to the 12 scenarios with asymmetric rates in different degrees. The *x*-axis shows the degree of asymmetry (RRD) of each CES rate. The dashed line at 0.05, indicates the expected *ED_95_* for the null model.

Because only asymmetry in the cladogenesis rate has a substantial effect, we zoom in on the comparison of the results of the symmetric scenario with the three scenarios that have different asymmetry levels in cladogenesis rate. Large inference error in ΔNESRTT only occurs when the mean cladogenesis rate is greater than the anagenesis rate (Fig. S2), which leads to fewer non-endemic species and more endemic species at present on the island.

### Effect of diversity dependence and state-dependent transitions

Diversity-dependence and state-dependent transitions have negligible effect on the inference errors in ΔNESRTT and σ_CS_, except when cladogenesis rates are asymmetric between states (Fig. 3 and S3). Among the four transition types, with low or high, symmetric or asymmetric transitions, parameter sets with higher transition rate from high rate state to low rate state (“low *q*_12_ high *q*_21_”) cause larger error in σ_CS_ than in the reverse direction (“high *q*_12_ low *q*_21_”) (Fig. 3 and S3).

**Fig 3.**
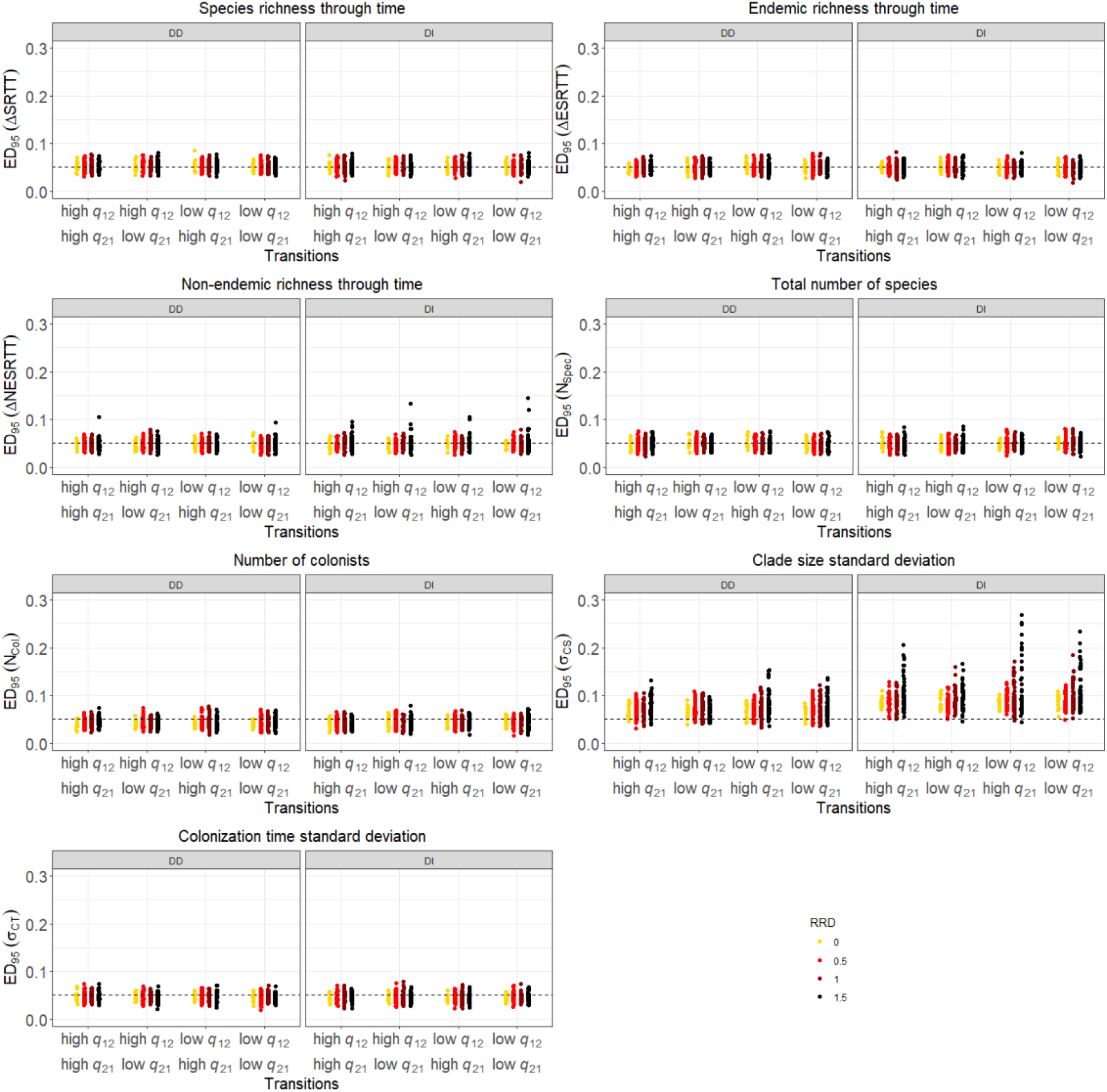
*ED_95_* of the seven metrics across all parameter combinations. The parameter sets are grouped by RRD, transition type and diversity-dependence. “DD” refers to diversity-dependent models that assume colonization and cladogenesis rates to be diversity-dependent, and “DI” refers to diversity-independent models that assume that all the CES rates are diversity-independent. The colors represent the asymmetry level (RRD) of CES rates between states. The dashed line at 0.05 indicates the expected *ED_95_* of the null model.

### Tip ratio and clade size variation

Large cladogenesis rate variation between states in SDS models can result in species richness variation between states, but it can also lead to clade size variation. State dependent transition rates may reinforce or reduce those variations to some extent. To understand what kind of empirical data may cause large errors using the DAISIE model, we checked the relationships between the inference errors with the tip ratio and the clade size variation in SDS results. We find that larger inference error does not always occur when the species richness is highly different between states, especially with higher rate transfer to lower rate state (“low *q*_12_ high *q*_21_”) (Fig. S4). In other words, tip ratio does not decisively control the robustness of the DAISIE model, because the phylogenies can be accurately reconstructed even with large richness differences between states (Fig. S4). The clade size variation of SDS results barely affects ΔNESRTT, but substantially affect the inference error in σ_CS_ (Fig. S5).

### Parameter estimation

When comparing the mean parameter values for simulations using the SDS model and the parameters inferred from the SDS results, the colonization and extinction rates are well estimated for most of the parameter sets (Fig. S6 and S7). However, nearly all scenarios show a systematic bias, with underestimated anagenesis and overestimated cladogenesis rates. The inference errors are positively correlated to the bias in cladogenesis and anagenesis when cladogenesis rates are largely asymmetric between states (Fig. S6 and S7).

## Discussion

Species traits are hypothesized to affect biological assemblages by altering diversification rates (Mitter et al., 1988; FitzJohn et al, 2009). Our results indicate that not incorporating the effect of trait dynamics on species diversification in the inference model and subsequently simulating the model with the obtained parameters allows surprisingly accurate reconstruction of the evolutionary history of species on an island under a wide range of scenarios. Hence, we conclude that the model is robust to leaving out the details of trait dynamics. Only in exceptional cases we see large differences between the simulations of a model with trait dynamics and a model without. This is specifically the case for two metrics: non-endemic richness through time (ΔNESRTT) and clade size standard deviation (σ_CS_).

Large differences between endemic species richness and non-endemic species richness may lead to large inference errors in ΔNESRTT. Within the parameter space investigated in this study, large error in ΔNESRTT occurs only when the mean cladogenesis rate between states is much higher than the mean anagenesis rate (Fig. S2). In this case, species with the higher cladogenesis rate state can rapidly speciate into a large clade, which leads to the endemic species richness being five to ten times the non-endemic species richness. Without accounting for trait dynamics, the estimated cladogenesis rate is closer to the higher cladogenesis rate than to the mean value of the two states in the SDS model. In contrast, the anagenesis rate is underestimated (Fig. S6 and S7). This leads to fewer non-endemic species, and more endemic species in the subsequent SIS simulations than in the SDS simulations, resulting in large error in ΔNESRTT.

The other metric whose estimation is affected by trait dynamics is σ_CS_. When trait states are conserved, and clades with a certain trait state have higher rates of diversification, clades with that trait state will likely become much more species-rich than clades with the other state. DAISIE assumes that all lineages diversify with the same rates, which generates balanced clades and leads to less clade size variation in the SIS results than in the SDS outputs. In addition, when cladogenetic speciation rate is diversity-dependent, competition between species in the same clade restricts the increase in species number, preventing clades from growing above a certain diversity level. Therefore, clade size cannot become extremely large, leading to lower error in σ_CS_ in diversity-dependent models than in diversity-independent models. However, we emphasize that even though DAISIE cannot accurately model the fine-scale variation between clades for some exceptional parameter combinations, it can still accurately reconstruct the dynamics of the whole community with multiple independent clades.

We attempted to determine the features of the data simulated under trait dependence that led to large inference errors in the two metrics where non-negligible error was found (ΔNESRTT, σ_CS_). The results indicate that clade size variation, which is the difference in species richness between clades, has a larger impact on the model accuracy than tip ratio, which is the species richness difference between states. This means DAISIE may cause error when fitting substantially unbalanced phylogenetic data, no matter if the variation between clades is caused by state-dependent diversification. In a study that used the DAISIE model to fit terrestrial birds of the Galapagos (Valente et al. 2015), the species richness of the clade of Darwin’s finches is reported to be much higher than the other clades of birds on the islands. The model that best fits the dataset assumes that Darwin’s finches have different cladogenesis and extinction rates than non-Darwin’s finches. Spectacular adaptive radiations such as Darwin’s finches are well-known on oceanic islands, and obviously lead to large clades. However, except for a handful of classic examples of adaptive radiations (Robichaux et al., 1990; Losos, 2009; Grant and Grant, 2008; Seehausen, 2006; Emerson, 2002), most island lineages do not diversify to form large clades (Patiño et al., 2017). Therefore, for most islands, clade size variation will rarely be extremely large when the whole assemblage of species of a given taxon is considered, suggesting the performance of models ignoring trait dynamics may not be affected for typical islands.

The power and accuracy of state-dependent biodiversity models has previously been evaluated with respect to the accuracy of the ancestral state reconstruction (Holland et al., 2020) and parameter estimation (Davis, et al., 2013). However, in these models the parameter inference model is based on the simulation model, which can generate simulated phylogenies for estimation. A statistical approach which does not rely on a formal model for coupling between states and diversification is available to detect the correlation between trait states and diversification with low type I error (Rabosky and Huang, 2016). However, this method cannot be used to reconstruct the ancestral state or to estimate parameters. In our approach, we apply a robustness analysis that uses the data from the complex model (SDS model) to evaluate the inference power of the simple model (SIS model). Complex models are vulnerable to overfitting of the data, and this leads to difficulty in accurately estimating parameters (Kelchner and Thomas, 2006). The pipeline used in this study identifies whether the simple model can accurately reconstruct diversity and phylogenies on islands without considering complex factors. In this way it constitutes a tool to determine whether it is useful to attempt to find a likelihood for the complex model or develop some other method to estimate parameters for the complex model. While we find that the trait-dependent DAISIE inference model seems to be robust to trait dynamics, it may still be meaningful to develop inference methods if one is interested in detecting the association between trait states and diversification, or in comparing diversification between mainland and island species with different traits (Patiño et al., 2017).

