## Supplementary for "A simple island biodiversity model is robust to trait dependence in diversification and colonization rates"

Supplementary Material

Supplementary Methods

In the SDS model, total colonization, diversification and transition rates of species with trait state *i* at time *t* are calculated as follows:

$$\lambda_{i}^{c}= \lambda_{0i}^{c}N_{i}(1- \frac{N}{K'})$$

$$\gamma_{i}= \gamma_{0i}M_{i}(1- \frac{N}{K'})$$

$$\mu_{i}= \mu_{0i}N_{i}$$

$$\lambda_{i}^{a}= \lambda_{0i}^{a}I_{i}$$

$$q_{ij}= q_{0ij}N_{i}$$

where *N_i_* is the present number of species on the island in trait state *i*; *M_i_* is the number of mainland species in state *i*; *N* is the total number of species present on the island (sum of all *N_i_*) descending from the same mainland species. *I_i_* is the number of non-endemic species with state *i* that are present on the island (i.e. those that have not undergone speciation). The parameters *λ_0i_^c^, µ_0i_, λ_0i_^a^, γ_0i_* are the per capita rates without diversity-dependence. *K’* is the maximum number of species in both states that can occupy in a clade. The cladogenetic and immigration rates are both state-dependent and diversity-dependent when *K’* is finite, and the function reduces to a diversity-independent version when *K’* = ∞. The transition rate from trait state *i* to state *j* is indicated by *q_ij_* with a per capita initial rate *q_0ij_*. The SDS model reduces to the SIS model when the CES rates of two states are the same.

Supplementary Figures

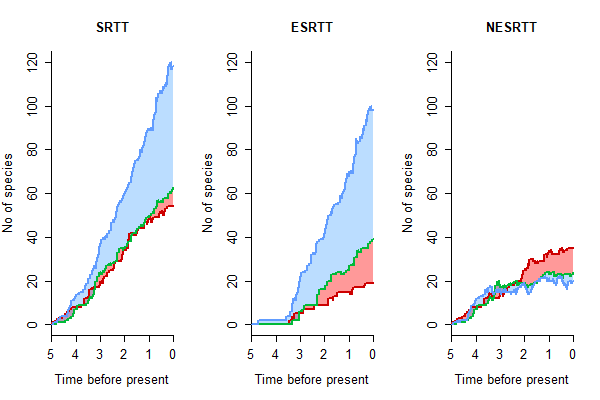

**Figure S1.** Illustration of nLTT (normalized lineage-through time) curves generated using the DAISIE model. The three panels show the three metrics: SRTT, ESRTT and NESRTT. The green, red and blue curves show the results of three replicate simulations. The area between two curves corresponds to the ΔnLTT (ΔSRTT, ΔESRTT or ΔNESRTT) of two simulations over time. The light blue area indicates the difference between the blue curve and the green curve, the red area indicates the difference between the red and the green curves.

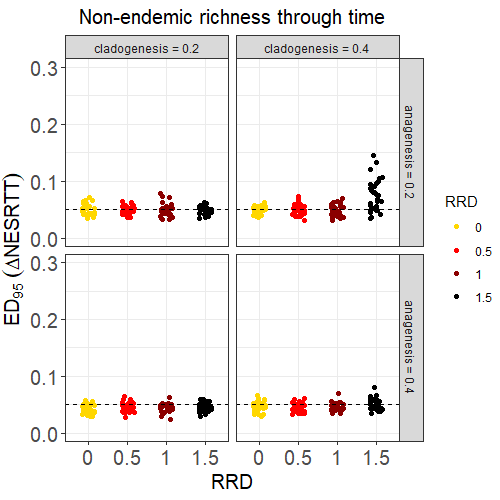

**Figure S2.** Distribution of *ED_95_* of ΔNESRTT across the scenarios with different asymmetry level in cladogenesis (scenarios 1-4 in Table 1C). Parameters are grouped into different combinations of mean cladogenesis and anagenesis rates (low and high rates). The colors represent the asymmetry level (RRD) of CES rates between states. The dashed line at 0.05, indicates the expected *ED_95_* for the null model.

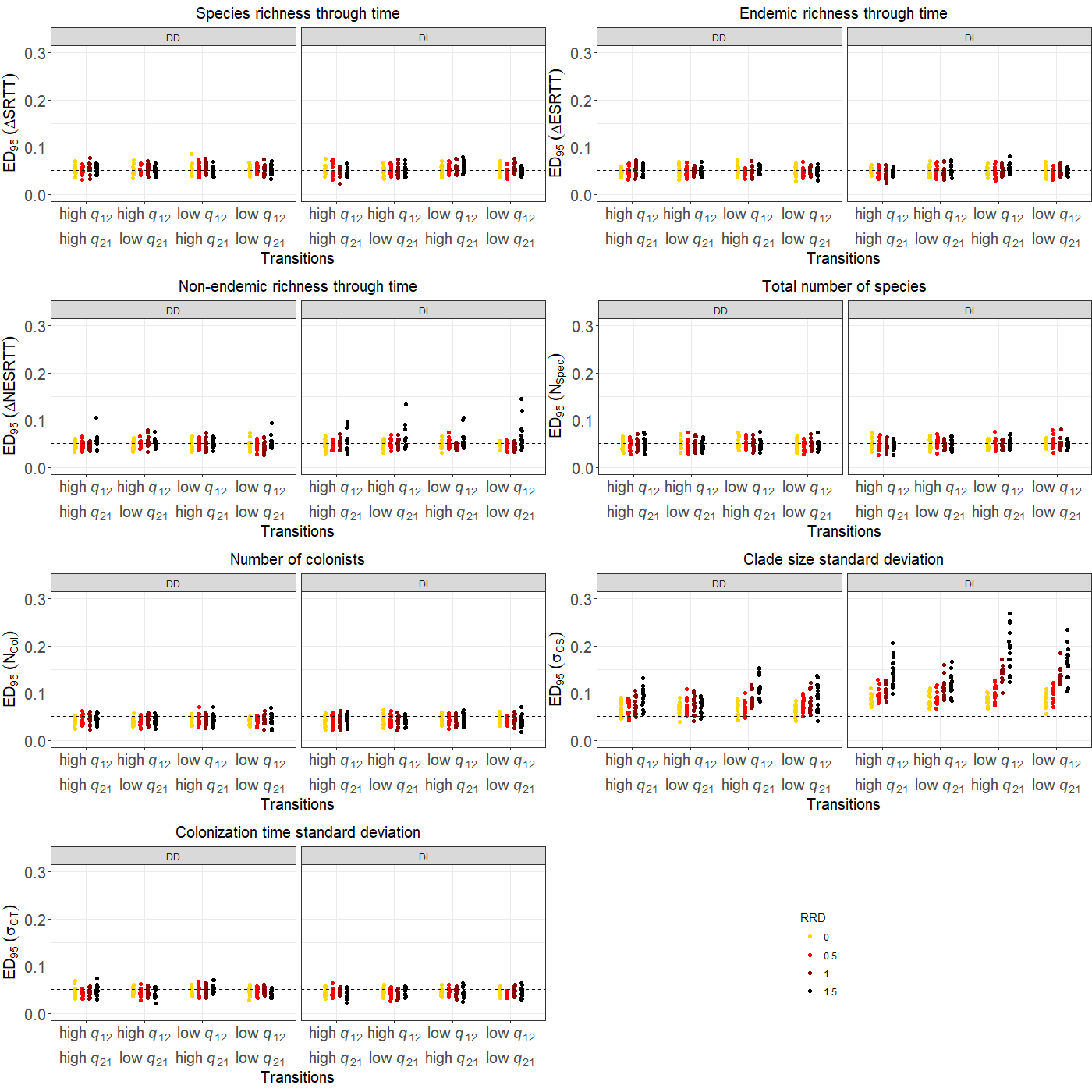

**Figure S3.** *ED_95_* of the seven metrics across the scenarios with different asymmetry level in cladogenesis (scenarios 1-4 in Table 1C). The parameter sets are grouped by RRD, transition type and diversity-dependence. “DD” refers to diversity-dependent model that considers colonization and cladogenesis rates are diversity dependent, and “DI” refers to diversity-independent model indicates all the CES rates are diversity independent. The colors represent the asymmetry level (RRD) of CES rates between states. The dashed line at 0.05 indicates the expected *ED_95_* of the null model.

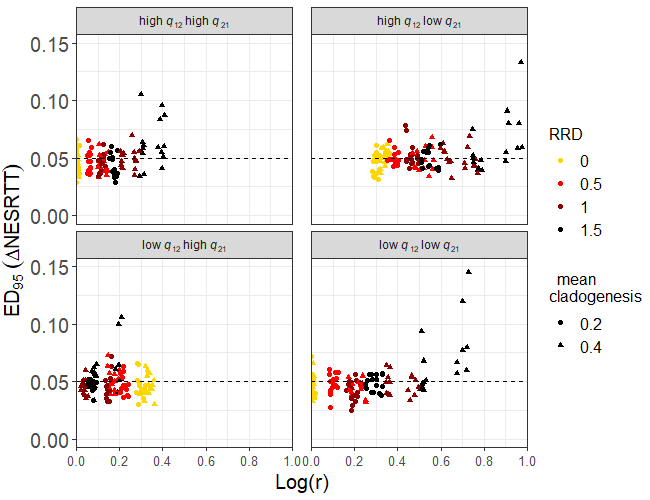

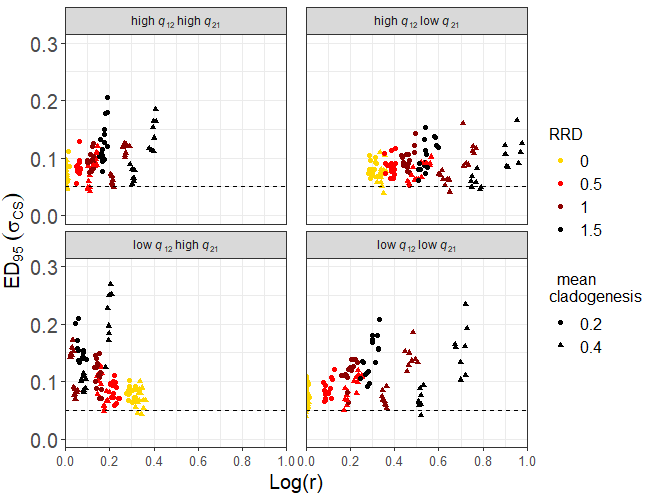

**Figure S4.** Relationship between tip ratio (r) with the *ED_95_* of ΔNESRTT and σ_CS_ across the scenarios with different asymmetry level in cladogenesis (scenarios 1-4 in Table 1C). The colors represent the asymmetry level (RRD) of CES rates between states. The shapes indicate different mean values of cladogenesis rate between states. The *x*-axis indicates the log value of the tip ratio. The dashed line at 0.05, indicates the expected *ED_95_* for the null model.

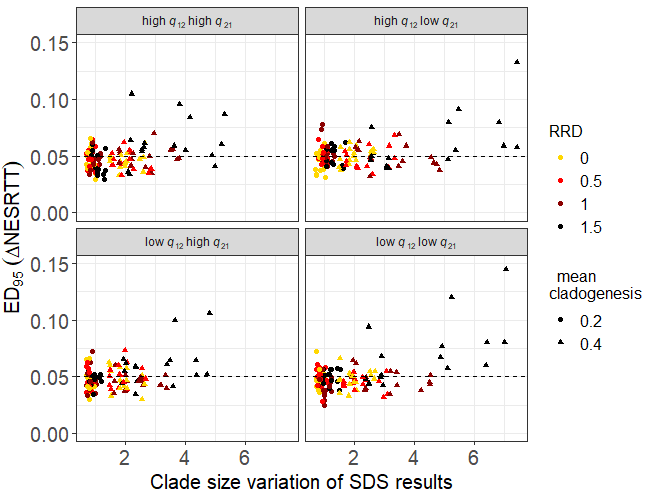

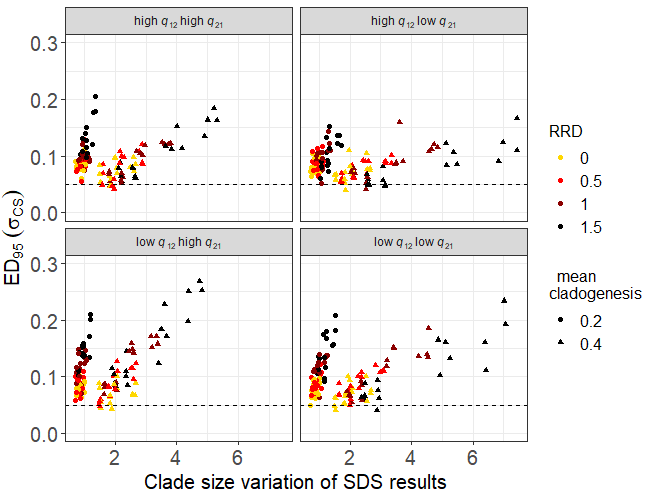

**Figure S5.** Relationship between the clade size variation of SDS results with the *ED_95_* of ΔNESRTT and σ_CS_ across the scenarios with different asymmetry level in cladogenesis (scenarios 1-4 in Table 1C). The colors represent the asymmetry level (RRD) of CES rates between states. The shapes indicate different mean values of cladogenesis rate between states. The dashed line at 0.05, indicates the expected *ED_95_* for the null model.

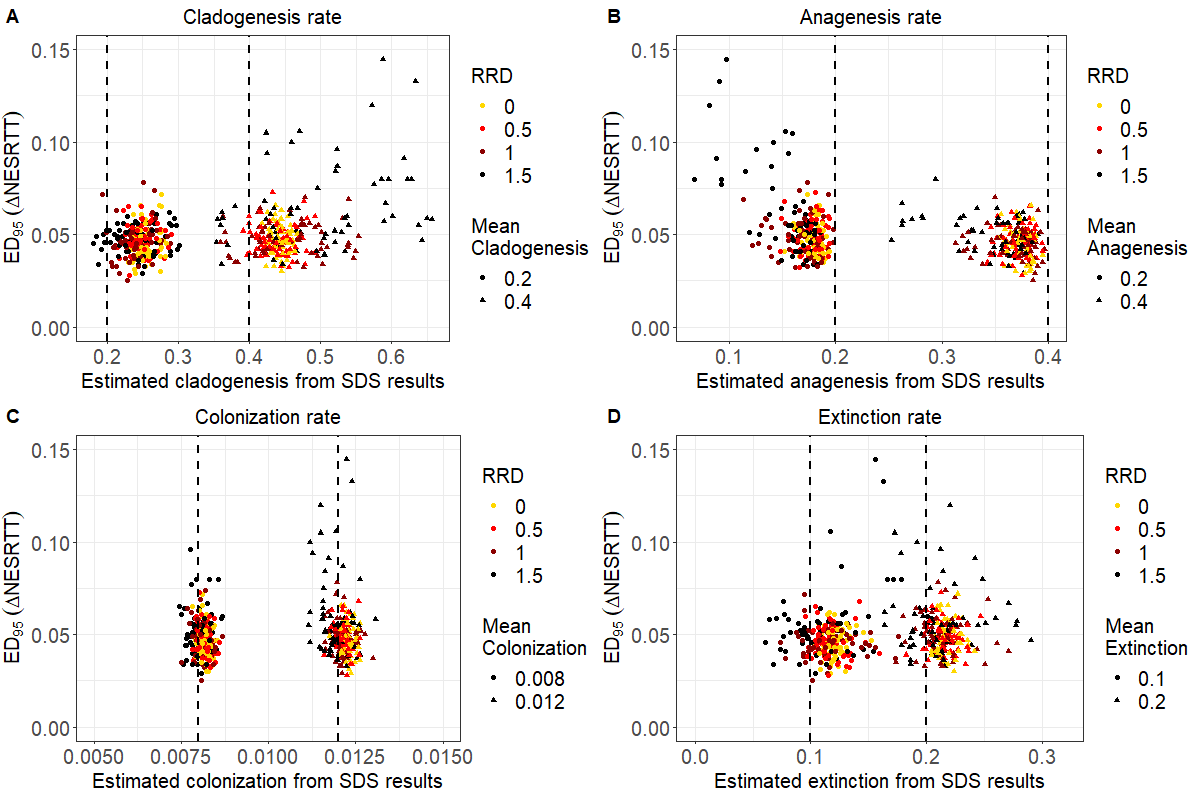

**Figure S6.** Relationships between the *ED_95_* of ΔNESRTT and the estimation bias across the scenarios with different asymmetry level in cladogenesis (scenarios 1-4 in Table 1C). Faceted plots show the estimation of (A) cladogenetic speciation rates, (B) anagenetic speciation rates, (C) colonization rates, (D) extinction rates from SDS results. The colors represent the asymmetry level (RRD) of CES rates between states. The shapes indicate different mean rate values between states. The dashed lines indicate the mean rate values in SDS models.

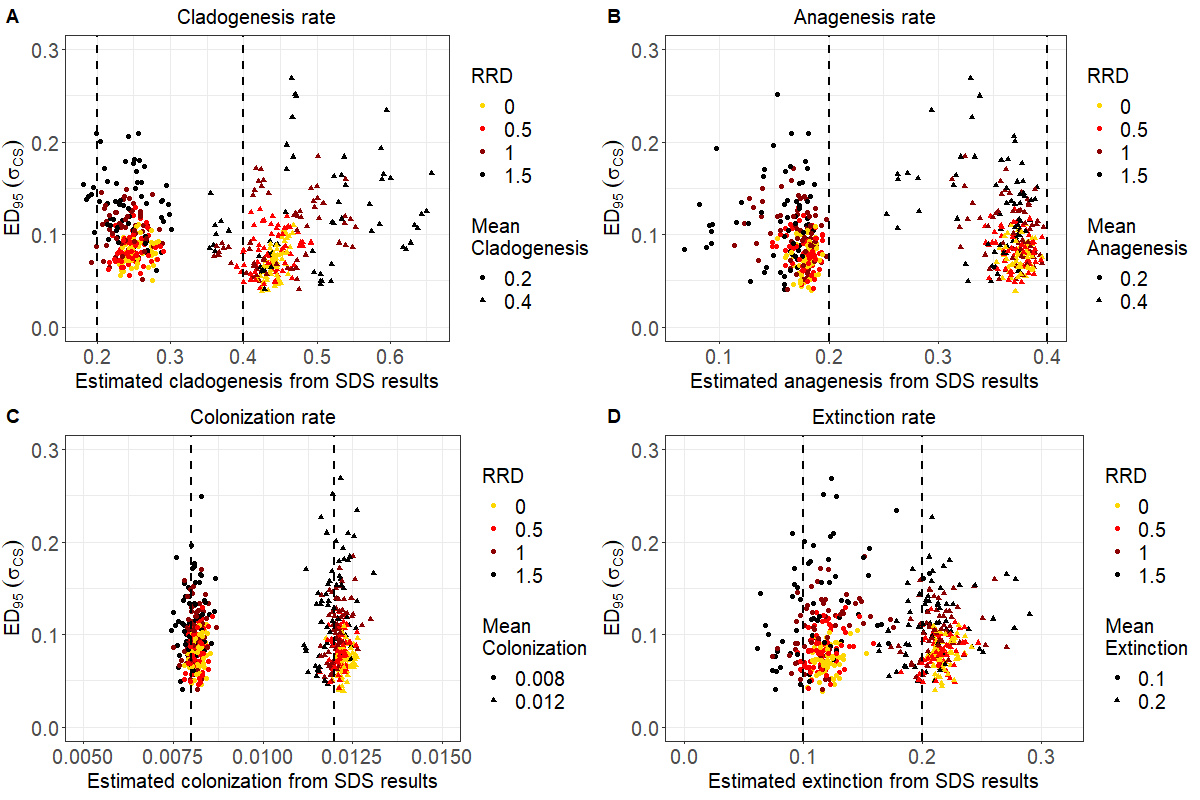

**Figure S7.** Relationships between the *ED_95_* of σ_CS_ and the estimation bias across the scenarios with different asymmetry level in cladogenesis (scenarios 1-4 in Table 1C). Faceted plots show the estimation of (A) cladogenetic speciation rates, (B) anagenetic speciation rates, (C) colonization rates, (D) extinction rates from SDS results. The colors represent the asymmetry level (RRD) of CES rates between states. The shapes indicate different mean rate values between states. The dashed lines indicate the mean rate values in SDS models.
